# GenomoBase: A comprehensive resource for the family *Genomoviridae*

**DOI:** 10.64898/2025.12.11.693732

**Authors:** Yair Cárdenas-Conejo

## Abstract

The family *Genomoviridae* comprises circular single-stranded DNA viruses reported from fungi, plants, animals and environmental samples. Although metagenomics has accelerated their discovery, genomic sequences, annotations and metadata remain dispersed across repositories. Here we present GenomoBase (https://www.genomobase.org), a curated resource that integrates genomic, ecological and bibliographic data for all 237 ICTV-recognized genomovirus species. GenomoBase incorporates Serratus-filtered SRA screening outputs, enabling prioritization of metagenomes for targeted re-analysis. As a proof of concept, a targeted bait-and-assemble workflow of one prioritized SRA run recovered two complete genomes, both below the 78% species demarcation threshold for genomoviruses. GenomoBase streamlines comparative analyses and metagenome-guided genome recovery.

---

High-throughput sequencing has transformed virus discovery, particularly for circular replication-associated protein-encoding single-stranded DNA (CRESS DNA) viruses. Within the phylum Cressdnaviricota, the family *Genomoviridae* has emerged as a globally distributed and ecologically diverse group. Since the initial description of *Sclerotinia sclerotiorum* hypovirulence-associated DNA virus 1 (SsHADV-1) [1], the family *Genomoviridae* has expanded to ten genera and 237 species recognized by the International Committee on Taxonomy of Viruses (ICTV) [2, 3]. Genomovirus genomes are small (∼1.9–2.4 kb) and typically encode a rolling-circle replication initiation protein (Rep) and a capsid protein (CP) in divergent orientation [4]. The Rep shares conserved rolling-circle replication motifs with geminivirus Reps, and the origin of replication commonly contains a stem–loop structure with a characteristic nonanucleotide motif (TAATATTAT), supporting rolling-circle replication [4]. In contrast, the CP forms a non-enveloped, T=1 icosahedral capsid and shows little recognizable similarity to capsid proteins of other virus groups [3]. Additional small open reading frames of unknown function are sometimes present.

Members of the family *Genomoviridae* have been reported from fungi, plants, animals and diverse environmental matrices, and sporadically from human-associated samples [5–9]. However, genomovirus genomes, annotations, host and geographic metadata, and the underlying metagenomic evidence remain scattered across general repositories and individual studies, complicating systematic comparison and dataset prioritization. To address this, we developed GenomoBase, a curated resource that consolidates ICTV-recognized reference genomes and harmonized metadata and incorporates Serratus-filtered Sequence Read Archive (SRA) screening outputs to support targeted mining and downstream genome recovery.

GenomoBase (https://www.genomobase.org) aggregates data for all 237 species currently recognized by the ICTV in the family *Genomoviridae* [2]. Reference genome records were retrieved from GenBank using the Virus Metadata Resource (VMR; VMR_MSL40.v2.20251013) as the source list. Using custom Python scripts (Biopython v1.86 [10]), we downloaded complete genome sequences and parsed CDS annotations (Rep and CP) together with associated metadata. Host and geographic fields were manually curated to improve consistency across records: host names were standardized and assigned to 11 ecological categories (mammals, birds, plants, insects, fungi, molluscs, reptiles, arachnids, actinopterygian fishes, humans and environmental samples), and free-text geographic descriptors were harmonized to country-level identifiers. The curated master table used to populate the database is provided as **Supplementary Table 1**.

The web resource was implemented in Django (Python v3.10+) with a PostgreSQL backend. Sequence similarity queries are supported through a local BLAST service (NCBI BLAST+ v2.17.0 [11]). An automated literature module periodically retrieves and categorizes recent genomovirus-related publications from PubMed.

To incorporate metagenomic evidence, we developed an “Explore Metagenomes” module based on Serratus screening outputs [12], which provide large-scale read-to-reference matches against genomovirus sequences. Serratus was queried in December 2025; at the time of querying, 660 SRA runs contained reads matching genomovirus references (alignment score ≥70; nucleotide identity ≥75%). In addition, 18 SRA runs were manually added based on the NCBI query “Genomoviridae” to include datasets with previously reported genomovirus signals not captured by the automated screen. Run-level metadata were retrieved via the NCBI and ENA APIs using a custom pipeline that harmonizes heterogeneous attributes, prioritizing host/isolation source and geographic information and parsing embedded coordinates when present. The resulting metagenome metadata file (**Supplementary File 1, JSON**) is downloadable from the portal and is used by the web application to power the Explore Metagenomes interface.

Analysis of the curated dataset indicates that the family is currently dominated by the genera *Gemycircularvirus* (126 species) and *Gemykibivirus* (50 species) (**Fig. 1A**). Genome lengths span ∼1.99–3.9 kb (mean ∼2.19 kb), consistent with compact CRESS DNA virus genomes (**Fig. 1B**); the upper bound reflects the combined length of the three genomic segments of the tripartite *Gemytripvirus fugra1*. Based on curated host assignments, mammals represent the largest category of unique species–host pairs (38%), followed by plants (15%) and birds (13.5%) (**Fig. 1C**). In the set of Serratus-retrieved SRA runs presented in Explore Metagenomes, environmental metagenomes are the most frequent sample type (26%), followed by human-associated (17%) and avian (13.6%) metagenomes (**Fig. 1D**). Using available location metadata, genomovirus-related read signals are detected in datasets from at least 44 countries across all continents, supporting a broad global distribution (**Fig. 1E**). The module offers a filterable table of Serratus-derived SRA runs annotated with alignment metrics (score and percent identity) with inferred location, host, and sample type, facilitating rapid triage and prioritization of public metagenomes for downstream genome reconstruction and comparative analyses.

**Figure 1.**
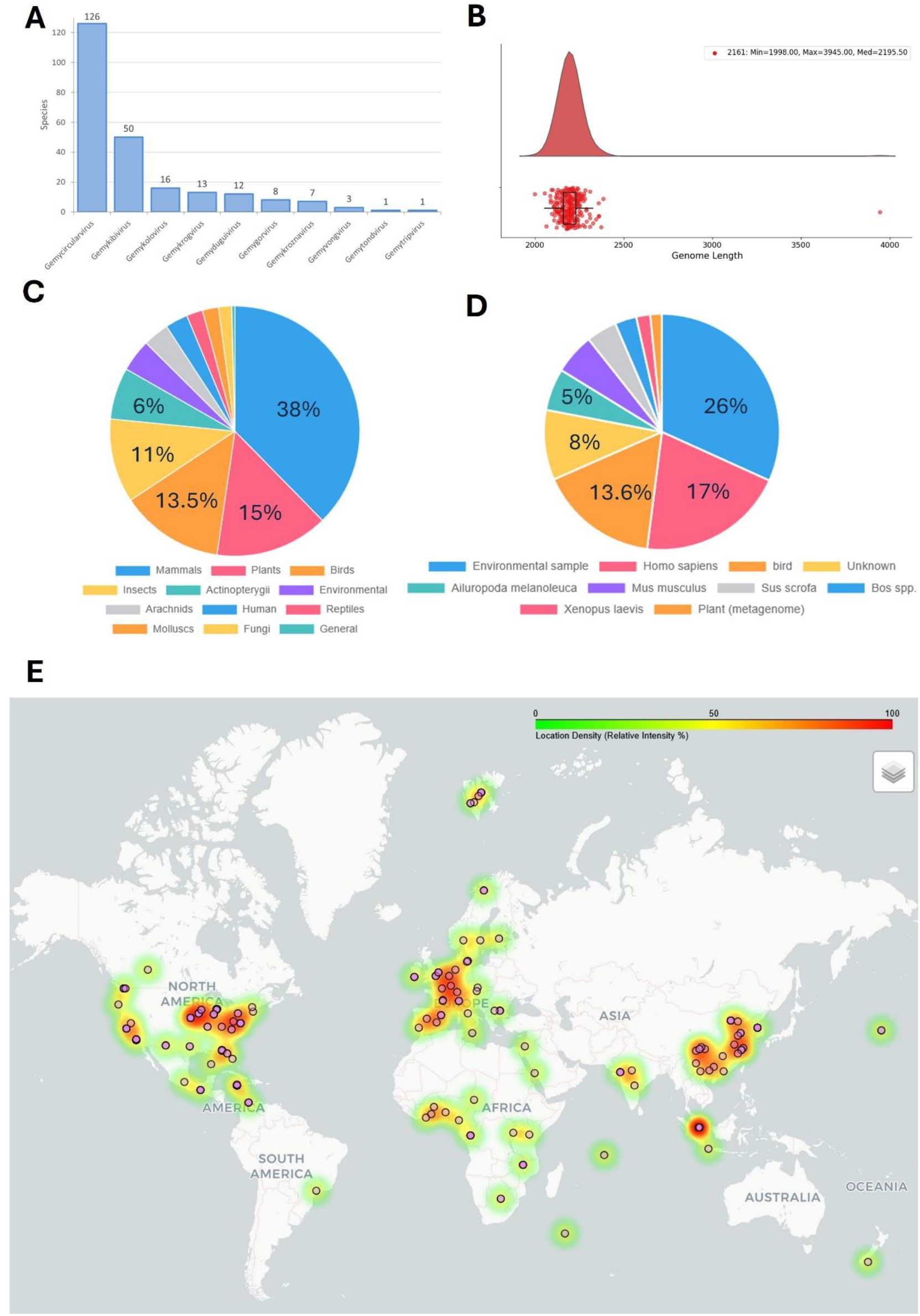
Overview of genomovirus species and metagenomic runs in GenomoBase. (A) Bar chart showing the distribution of currently classified species across ICTV-recognized genera. (B) Genome length distribution visualized as a raincloud plot combining a density curve, individual genomes plotted as points, and a boxplot. (C) Broad host categories for genomoviruses, summarized as a pie chart based on curated host metadata. (D) Pie chart summarizing SRA runs with genomovirus hits, grouped into broad host or sample categories (e.g. birds, humans, domestic pigs, environmental and others). (E) Global heat map of sampling locations for the 282 runs with latitude/longitude metadata, illustrating the relative density of genomovirus-positive metagenomes. Warmer colours indicate higher local density across countries and regions.

To demonstrate the utility of GenomoBase for prioritized data mining and taxonomic contextualization, we used the Explore Metagenomes module to select a public dataset for downstream local analysis. Using the built-in filters, we restricted the Serratus-derived SRA runs to those with an alignment score of 100 and a percent identity of 80–90% to prioritize divergent candidates for genomovirus genome recovery. We selected the SRA run SRR4201716 (BioSample SAMN05725476), derived from urban bat (*Molossus molossus*) saliva collected in French Guiana [13], because it showed the lowest percent identity (83%). Raw reads were downloaded and subjected to a targeted “bait-and-assemble” workflow. Briefly, the complete GenomoBase reference set (n=237) was used as bait to recruit reads with BBMap v38.90 (Bushnell, BBMap; https://sourceforge.net/projects/bbmap/) using a permissive identity threshold (minid=0.70) to capture divergent lineages, and recruited reads were assembled de novo with SPAdes v3.15.5 [14] using the --careful option.

This downstream workflow yielded 15 contigs (**Supplementary File 2, FASTA**), of which two were manually validated as complete, circular genomovirus genomes and were selected for downstream analyses by confirming consistent read mapping across the entire sequence (including reads spanning the circularization junction), the absence of ambiguous bases (Ns), and no evidence of chimeric assembly. Other contigs corresponded to partial assemblies and/or showed high similarity to known genomes and were not pursued further.

The first genome, designated “Genomoviridae SRR4201716 Contig 1” (**GenBank accession pending;** BankIt 3033975), is 2,197 bp in length. Structural inspection identified the genomovirus-typical stem–loop origin of replication containing the nonanucleotide motif TAATATTAT. The genome encodes two major open reading frames in divergent orientation (Rep and CP), as well as a putative ORF3 (**Fig. 2A**). A total of 8,961 reads aligned to the assembled genome, yielding 100% coverage across the full length (**Fig. 2C**). BLASTn searches against GenomoBase identified the closest classified relative as *Gemycircularvirus echiam1* (KY308268.1) with 87% nucleotide identity over 27% query coverage (E-value = 0.0), consistent with a partial local alignment rather than genome-wide similarity.

**Figure 2.**
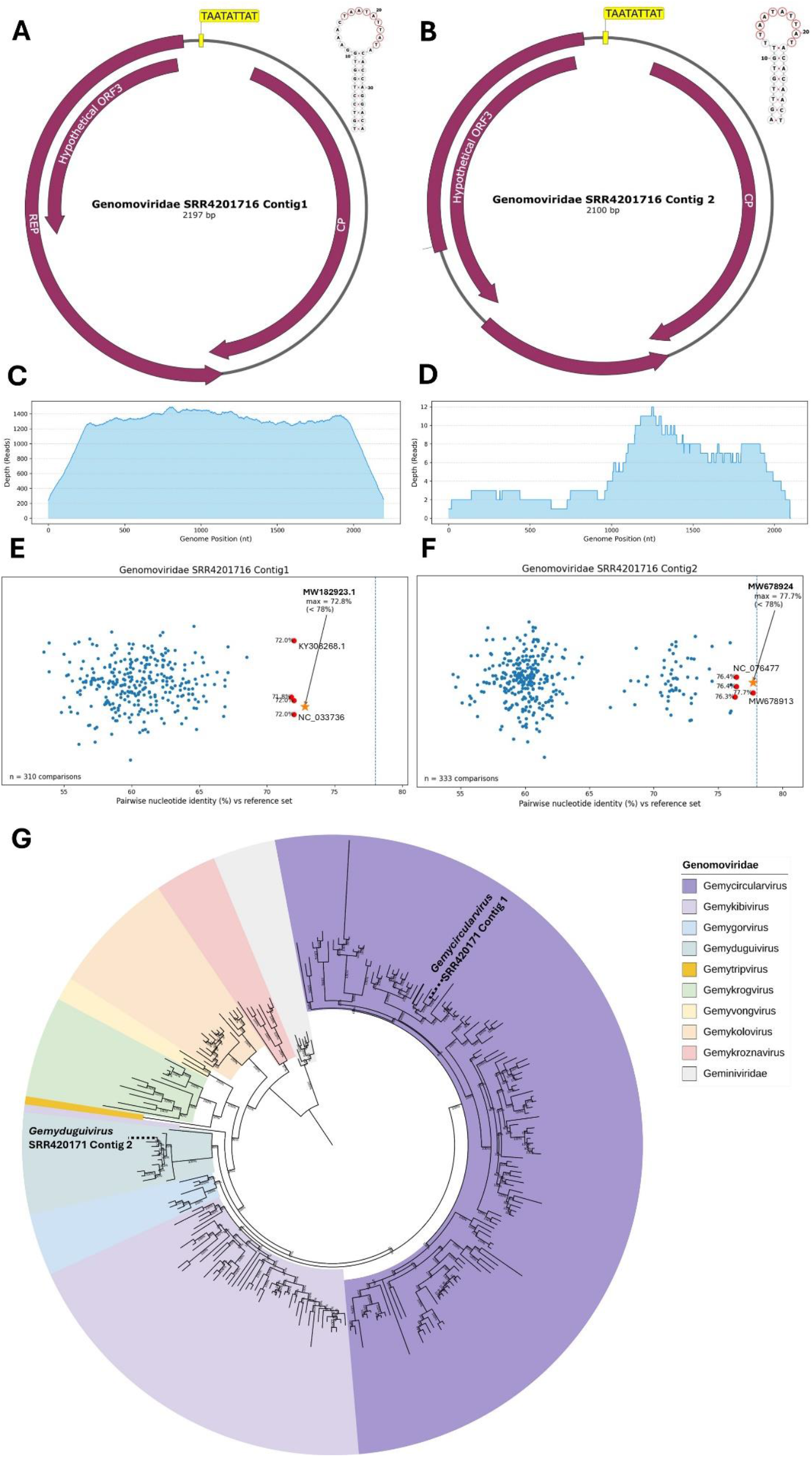
Genomic organization, read support, nucleotide identity, and phylogenetic placement of two genomovirus genomes assembled from metagenomic dataset SRR4201716. (A–B) Circular genome maps of Genomoviridae SRR4201716 Contig 1 (2,197 bp) and Genomoviridae SRR4201716 Contig 2 (2,100 bp), showing annotated ORFs (Rep, CP, and a hypothetical ORF3) and the conserved origin-of-replication motif in the intergenic region (highlighted in yellow), together with the predicted stem–loop structure. (C–D) Read depth profiles across the genome for Contig 1 (C) and Contig 2 (D) based on read mapping to each assembly (x-axis, genome position; y-axis, read depth). (E–F) Whole-genome pairwise nucleotide identity spectra for Contig 1 (E) and Contig 2 (F) relative to complete genomes in GenomoBase and the top BLAST matches; each point represents a single pairwise comparison (n = 310 and n = 333, respectively). The dashed blue vertical line denotes the species demarcation threshold (78% identity). The highest identity was 72.8% for Contig 1 and 77.7% for Contig 2 (both <78%), highlighted by a yellow star; the remaining top five matches are highlighted in red. (G) Rep protein phylogenetic analysis used to infer genus-level placement of the putative novel viruses. Rep sequences were aligned with MAFFT v7 [17] and trimmed with trimAl [18]. A maximum-likelihood tree was inferred with IQ-TREE v3 [16], and the figure was visualized and edited in iTOL [19]. The study contigs are labeled in the tree. The full phylogeny is provided in Supplementary Figure S1.

The second complete circular genome (“Genomoviridae SRR4201716 Contig 2”; **GenBank accession pending**; BankIt 3033975) is 2,100 nt and contains the TAATATTAT nonanucleotide at the predicted stem-loop origin. The genome encodes CP and Rep, and a predicted overlapping RepA-like ORF, as well as a putative ORF3 (**Fig. 2B**). A total of 35 reads mapped to the assembled genome, providing 99.8% coverage (**Fig. 2D**); importantly, mapping supported the circularization junction, and no ambiguous bases (Ns) were present. BLASTn searches identified the closest classified relative as *Gemyduguivirus arteca1* (MN823676.1) with 93.6% identity over 30% query coverage (E-value = 0.0). Despite the low read depth, the near-complete coverage together with read support across the circularization junction supports the integrity of the assembly.

To assess whether these genomes represent novel species, we applied the species demarcation criterion proposed by Varsani and Krupovic [4]. Genome-wide pairwise nucleotide identities were calculated with SDT v1.3 [15] using alignments that included the complete genomes of the 237 ICTV-recognized genomovirus species represented in GenomoBase, together with the top BLASTn hits retrieved from GenBank. In contrast to BLASTn, which reflects local similarity, SDT reports genome-wide pairwise identity used for species demarcation. In both cases, the maximum pairwise identity was below the 78% threshold (**Fig. 2E,F**; **Supplementary Table 2**), consistent with assignment as candidate distinct species under this criterion. Specifically, “Genomoviridae SRR4201716 Contig 1” showed a highest identity of 72.8% to Gemycircularvirus isolate bfb129gen4 (MW182923.1), whereas Contig 2 showed a highest identity of 77.7% to Genomoviridae sp. isolateD9459 (MW678924.1). To corroborate genus-level placement, we inferred a maximum-likelihood phylogeny of the Rep protein in IQ-TREE v3.0.1[16] (1,000 ultrafast bootstrap replicates) using an alignment comprising the 237 GenomoBase Rep sequences; this analysis placed Contig 1 within *Gemycircularvirus* and Contig 2 within *Gemyduguivirus* (**Fig. 2G**). Together, these results illustrate how GenomoBase can guide the selection of informative public metagenomic datasets and support the recovery and taxonomic contextualization of complete genomes of the family *Genomoviridae* from SRA data.

In conclusion, GenomoBase provides a centralized and curated resource for the family *Genomoviridae* by integrating ICTV-based reference genomes with standardized host and geographic metadata, an automatically updated literature catalogue and built-in tools for sequence similarity searches. In addition, the Explore Metagenomes module incorporates SRA screening outputs into a filterable and downloadable interface, allowing users to move efficiently from curated references to candidate public datasets for downstream genome reconstruction and comparative analyses. As demonstrated by our proof-of-concept analysis, this workflow can support the prioritization of divergent runs and the recovery of complete circular genomes, followed by genome-wide identity and phylogenetic placement within an ICTV-based framework. By lowering the practical barriers to comparative analyses and targeted mining of public metagenomes, GenomoBase is expected to facilitate ecological and evolutionary studies of the family *Genomoviridae* and to accelerate the identification of under-sampled host and environmental contexts. The resource will be updated to reflect future ICTV releases and the continued expansion of publicly available metagenomic data.

## Acknowledgements

This study was supported by grants from the Secretaría de Ciencia, Humanidades, Tecnología e Innovación (CBF-2025-G-902).

## Statements and Declarations

### Authors’ contributions

Y.C.-C. conceived and designed the study, developed the database and web server, curated and harmonized the data, performed the analyses, generated figures and tables, and wrote and revised the manuscript.

### Data availability

The online instance of GenomoBase is freely accessible without registration at https://www.genomobase.org. Curated datasets, processed tables, and the custom scripts used to build and update GenomoBase, together with sample input data and corresponding output, are publicly available via Zenodo (DOI: 10.5281/zenodo.17860246). The two complete circular genomes reported here have been submitted to GenBank (BankIt submission 3033975; accession numbers pending). For peer review, both sequences are provided as the BankIt GenBank-format submission file uploaded with the manuscript (BankIt 3033975; accessions pending).

