## Supplementary figures and images for "GenomoBase: A comprehensive resource for the family *Genomoviridae*"

### Supplementary Figure 1

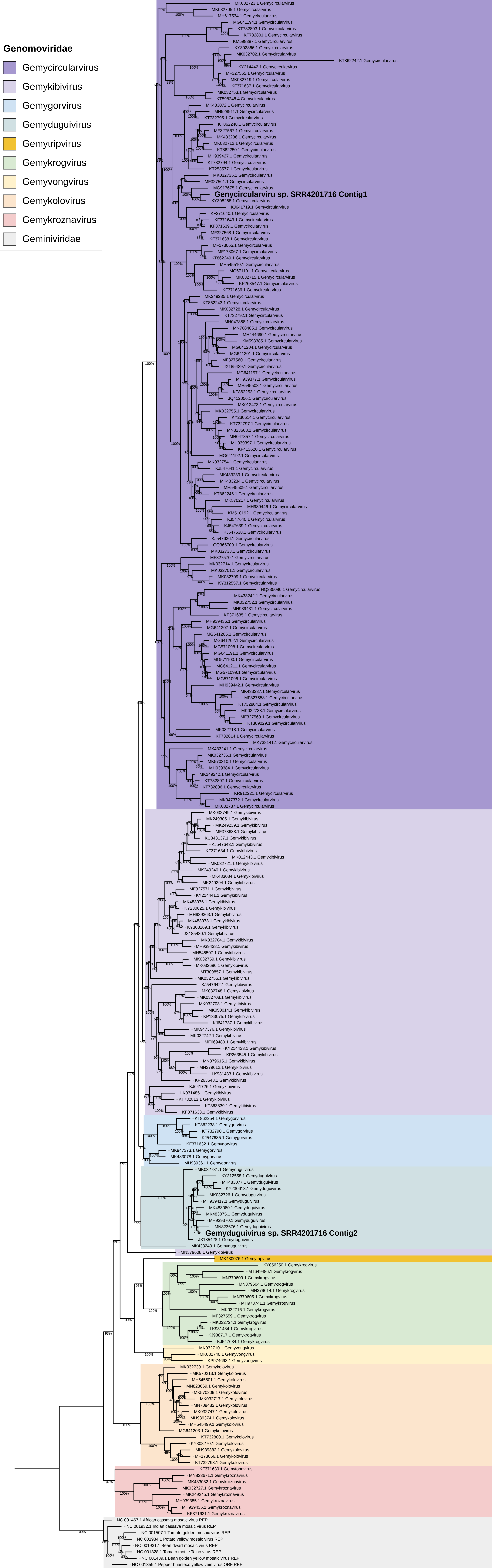
